# Lean breast adipocytes secrete an oxylipin that suppresses breast cancer via ferroptosis

**DOI:** 10.1101/2025.07.17.665258

**Authors:** Meghan C. Curtin, Abigail E. Jackson, Elisabeth A. Brown, J. Alan Maschek, David Lum, James E. Cox, Alana L. Welm, Keren I. Hilgendorf

## Abstract

Obesity is predicted to become the largest modifiable risk factor for breast cancer in postmenopausal women, yet the mechanisms underlying this association are unclear. We identified a novel role for the endogenous oxylipin 9S-HODE, secreted by lean adipocytes, to induce ferroptosis in breast cancer cells while sparing normal breast epithelial cells. Obese adipocytes fail to secrete 9S-HODE, suggesting that the loss of ferroptosis induction significantly contributes to the acceleration of obesity-associated breast cancer. Consequently, the inhibition of ferroptosis accelerates breast cancer in lean, but not obese, mice. Further, 9S-HODE inhibits the growth of patient-derived breast cancer organoids, and supplementing 9S-HODE into tumors in obese mice is sufficient to reduce tumor burden, underscoring its potential as a therapeutic agent.

## Main Text

Breast cancer affects 1 in 8 women, resulting in over 40,000 deaths each year in the US (*1, 2*). The risk of developing post-menopausal breast cancer increases dramatically with obesity, with each 5-unit increase in body mass index (BMI) corresponding to a 12% increase in risk (*3*).

Regardless of menopausal status, obesity also increases the risk of large, high-grade tumors (*4-7*). As 70% of US adults are overweight or with obesity and obesity rates are expected to continue rising (*8, 9*), it is critical to understand the mechanisms underlying breast cancer acceleration in obesity and provide targeted therapeutics to the significant fraction of breast cancer patients with obesity.

The transcriptome, lipidome, and secretome of adipocytes within breast tissue are substantially altered with obesity (*10-18*), and several groups have demonstrated how cancer-associated adipocytes in obese mice and patients can promote breast cancer (*19, 20*). For example, Maguire et al. showed that adipocytes in obese mice promote breast cancer progression by releasing creatine (*21*). Obesity also promotes the increased secretion of adipokines, including resistin and leptin, and miRNAs, such as miR-155, which enhance the proliferation and epithelial-to-mesenchymal transition of breast cancer cells (*22-25*). Finally, increased aromatase activity linked to increased adiposity can substantially elevate estrogen levels and promote the growth of estrogen-driven breast cancer in postmenopausal women with obesity (*26*). Beyond proteins, metabolites, and miRNAs, adipocytes also secrete lipids, although the role of lipid signaling in breast cancer remains underexplored. Studies have shown that obesity alters both serum and adipose tissue lipid profiles, and that the established lipid signatures linked to metabolic health are highly conserved between mice and humans (*18*). In particular oxylipins, which are oxidized polyunsaturated fatty acid (PUFA) derivatives, have been identified as bioactive lipid signals that play a significant role in cardiovascular disease, inflammation, neurological disorders, and cancer (*27*). Several oxylipins have been shown to trigger ferroptosis, a form of cell death in which iron-containing enzymes or iron itself react with oxygen and PUFAs to generate high levels of lipid peroxides in cellular membranes, resulting in membrane rupture (*28-33*).

Here, we describe a potent ability of oxylipins secreted by lean mammary adipocytes to suppress breast cancer growth. We demonstrate that one of these oxylipins, 9S-HODE, triggers ferroptosis in breast cancer cells. The absence of 9S-HODE in obese mice results in breast cancer cells undergoing minimal ferroptosis, leading to accelerated growth compared to lean mice.

Accordingly, inhibiting ferroptosis accelerates breast cancer in lean but not obese mice. Finally, exogenous administration of 9S-HODE into tumors in obese mice is sufficient to slow tumor growth compared to controls, and 9S-HODE also inhibits the growth of patient-derived breast cancer organoid cultures. Together, these data support a model in which breast cancer is accelerated in obesity at least in part due to the loss of tumor-suppressive lipid secretion by obese adipocytes, and restoring this lipid has therapeutic potential.

## Results

### Lean adipocytes suppress breast cancer cell growth

Previous studies have demonstrated that E0771 breast cancer cells injected orthotopically into obese C57BL/6 mice form significantly larger tumors when compared to lean mice (*21*). To confirm and expand on these observations, female C57BL/6J mice were either placed on a high fat diet (HFD) or a normal chow diet for 9 weeks. Mice on a normal chow diet have lower body weight and lower fat mass than the mice on HFD, and will henceforth be referred to as lean, versus obese, mice (fig. S1A-B). E0771 cells were injected into the mammary adipose tissue (commonly referred to as the mammary fat pad) of lean or obese mice. Consistent with previous studies, E0771 tumor growth was significantly accelerated in obese mice relative to lean mice (Fig. 1A). Similarly, tumor growth of another syngeneic breast cancer cell line, Py230, was accelerated in obese mice relative to lean mice (fig. S1C-D). However, the absence of a default tumor growth rate for reference precludes our ability to conclude if tumor growth is suppressed in lean mice and/or accelerated in obese mice.

**Fig. 1.**
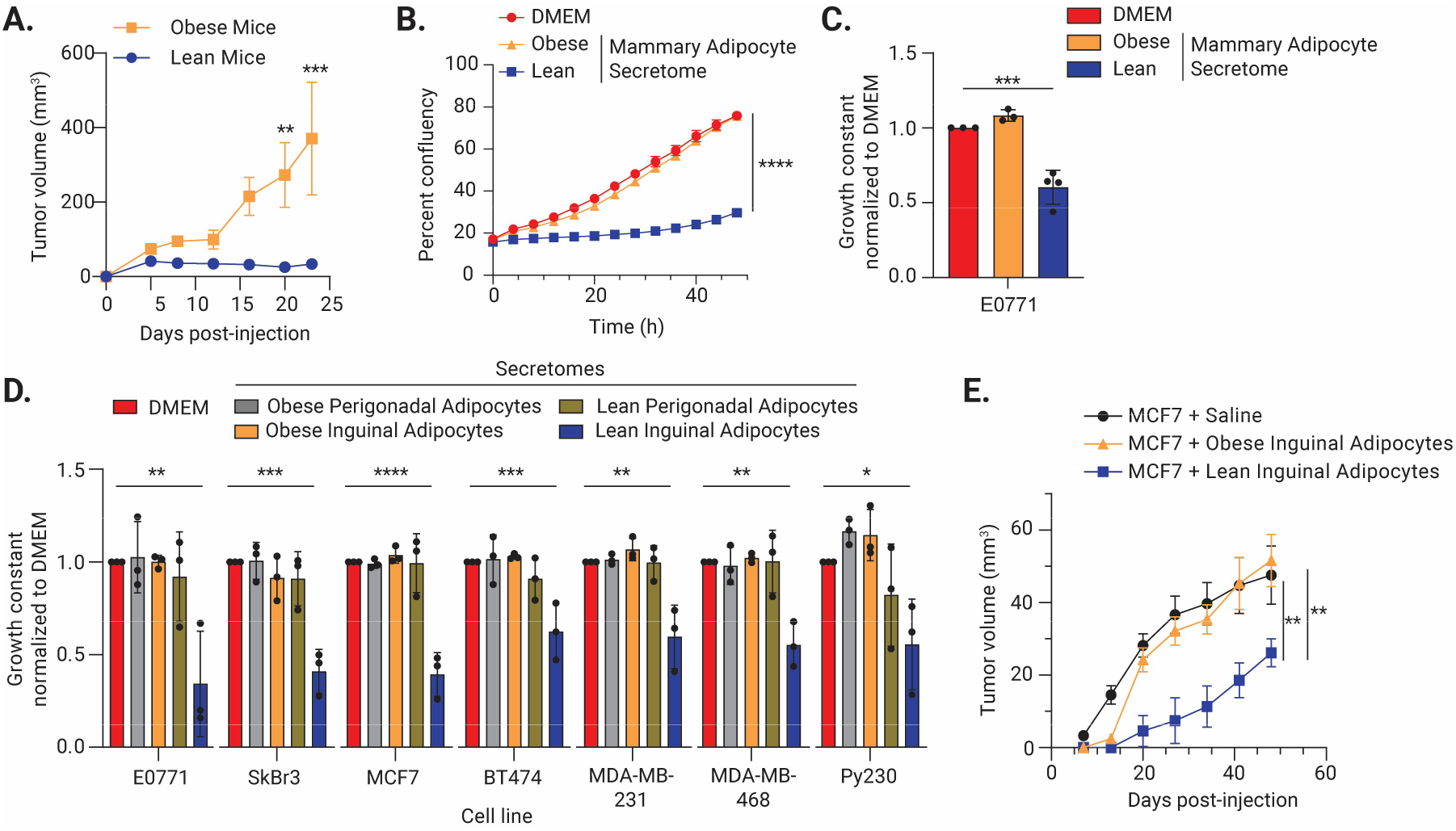
Lean inguinal adipocytes inhibit breast cancer cell growth. (**A**) Orthotopic growth of E0771 mammary tumors in C57BL/6J mice on a high fat (obese) diet or standard (lean) diet (n=10). **(B-C)** Lean, but not obese, mammary adipocytes suppress breast cancer cell growth. (B) is a representative growth curve; (C) is the calculated growth constant for E0771 cells treated with the secretome of 3-4 individual isolations of mammary adipocytes (n=3-4). (**D**) The lean inguinal adipocyte secretome decreases the growth constant of seven breast cancer cell lines.

To differentiate between suppression of tumor growth in lean mice versus acceleration of growth in obese mice, we examined the effect of paracrine signals from isolated lean and obese adipocytes on breast cancer cell growth *in vitro*. To collect paracrine signals, mammary adipocytes from lean or obese female mice were isolated and placed in culture. Consistent with prior characterization (*34*), isolated lean adipocytes were smaller than obese adipocytes (fig. S1E). The media containing the factors secreted by lean or obese adipocytes was collected and will henceforth be referred to as the secretome. The lean or obese secretome was then supplemented with the standard culture media in a 1:1 ratio and added to proliferating E0771 cells *in vitro*. The growth constant of E0771 cells was calculated using the IncuCyte live-cell imaging system measuring changes in 2D confluency over time. Strikingly, E0771 growth was significantly decreased in the presence of the lean mammary secretome compared to normal growth media alone (Fig. 1B-C). There was no effect on E0771 growth in the presence of the obese mammary secretome.

To determine if this is a sex-specific effect, male C57BL/6J mice were placed on HFD or normal chow. Male mice on normal chow had lower body weight and fat mass compared to mice on a HFD (fig. S1F-G). Of note, the inguinal adipose tissue in female mice contains the mammary glands. Next, we isolated and cultured inguinal adipocytes from lean and obese male mice, then collected the secretome. We observed a comparable decrease in E0771 growth *in vitro* in the presence of either the male inguinal or female mammary secretome (Fig. 1B, fig. S1H). The obese male inguinal secretome had no effect on E0771 growth (fig. S1H). Because both male and female lean adipocytes significantly suppressed breast cancer cell growth, male inguinal adipose tissue was utilized for further mechanistic interrogation as it is larger than female adipose tissue, particularly in lean mice. Key data are confirmed with female mammary adipose tissue.

Strikingly, the secretomes of lean inguinal adipocytes robustly suppressed breast cancer cell growth across multiple human and murine breast cancer cell lines, representative of all clinical subtypes of breast cancer (Fig. 1D). No growth effect was observed with obese inguinal secretome or the secretome of adipocytes from the visceral perigonadal adipose depot of lean or obese male mice (Fig. 1D). The lean inguinal secretome’s suppression of breast cancer cell growth was not due to nutrient depletion in the secretome, as replacing the secretome with serum-free DMEM or HBSS in a 1:1 ratio with standard growth media had no effect on E0771 or SkBr3 growth (fig. S1I). Thus, lean, but not obese, inguinal/mammary adipocytes secrete an inhibitory factor(s) that robustly suppresses breast cancer cell growth *in vitro*.

To confirm this effect *in vivo*, MCF7 breast cancer cells were combined with adipocytes freshly isolated from the inguinal adipose tissue of lean or obese male C57BL/6J mice or a saline control. The mixtures were injected subcutaneously into the flank of immunocompromised NRG mice, and tumor growth was monitored over time. Lean adipocytes robustly inhibited MCF7 growth *in vivo*, while we observed no difference in tumor growth with the co-injection of obese adipocytes compared to saline control (Fig. 1E). Thus, lean adipocytes are sufficient to suppress breast cancer cell growth *in vivo* and *ex vivo*.

### Lean adipocytes trigger ferroptosis in breast cancer cells

Lean inguinal adipocytes may suppress breast cancer cell growth via a combination of decreasing proliferation and/or increasing cell death. To establish the mechanism(s) of growth suppression, we next compared the gene expression of E0771 and SkBr3 cells treated with the lean secretome to those treated with standard growth media (Data S1, S2). This revealed an upregulation of transcripts linked to the ferroptosis pathway (Fig. 2A, fig. S2A-B). To confirm this, the levels of the reactive aldehyde 4-HNE, a byproduct of lipid peroxidation and marker of ferroptosis, were assessed (*28, 35*). E0771 cells had high levels of 4-HNE when treated with lean secretome compared to standard growth media and similar to the levels observed with the ferroptosis inducer ML-162 (Fig. 2B). To test if E0771 growth suppression by the lean adipocyte secretome was mediated by increased ferroptosis, cells were co-treated with ferroptosis inhibitors. E0771 growth was completely rescued in the presence of the ferroptosis inhibitor Ferrostatin-1 (Fig. 2C) or Liproxstatin-1 (fig. S2C). Finally, we confirmed that E0771 cells treated with the lean secretome were undergoing cell death (Fig. 2D). To determine the contribution of other cellular processes to growth suppression, we measured apoptosis using a fluorescent reporter of cleaved caspase 3/7 activity. E0771 and SkBr3 cells showed no increase in apoptosis in the presence of lean secretome (fig. S2D). We also tested if the lean secretome induced cell cycle arrest by quantifying propidium iodide incorporation and phospho-Histone H3 levels. The lean secretome induced an S-phase cell cycle arrest in both E0771 and SkBr3 breast cancer cells (fig. S2E-J). We do not know if the induction of ferroptosis and S-phase cell cycle arrest are mechanistically linked. Taken together, our results show that the lean adipocyte secretome suppresses E0771 growth by inducing ferroptotic cell death and S-phase cell cycle arrest.

**Fig. 2.**
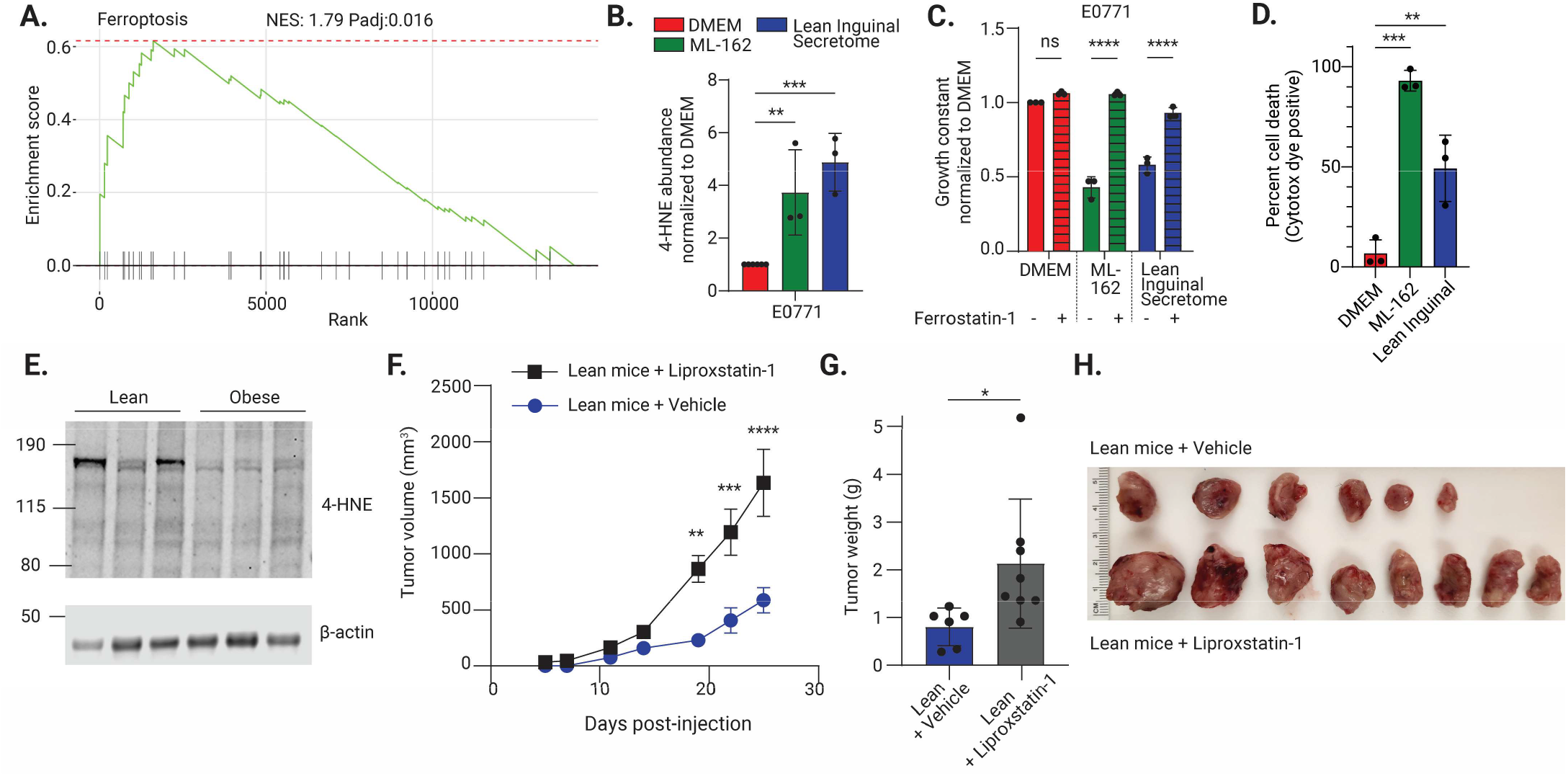
Lean inguinal adipocytes induce ferroptosis in breast cancer cells. (**A**) Gene set enrichment analysis for ferroptosis-related genes in E0771 cells treated with the lean inguinal adipocyte secretome compared to DMEM control. (**B**) E0771 cells treated with the lean inguinal adipocyte secretome or the ferroptosis inducer ML-162 have higher levels of 4-HNE than DMEM control (n=3). (**C**) Treatment with ferroptosis inhibitor ferrostatin-1 rescues the growth of E0771 cells treated with the lean inguinal adipocyte secretome or ML-162 (n=3). (**D**) Cytotox dye incorporation confirms that ML-162 and the lean inguinal adipocyte secretome induce cell death (n=3). (**E**) Immunoblot for 4-HNE in the tumor lysates of E0771 mammary tumors taken from lean or obese mice demonstrates an increase in 4-HNE abundance in tumors from lean mice (n=3 mice). (**F-H**) Growth of E0771 mammary tumors is (F) accelerated in lean mice with daily injections of liproxstatin-1 (n=8) compared to vehicle control (n=6). (G) Tumor weight and (H) images taken at endpoint. (B-D, G) Data are mean ± SD, (F) data are mean ± SEM. p-values calculated using one-way ANOVA followed by Dunnett’s multiple comparison test (B, D) or Šídák’s multiple comparisons test (C), two-way ANOVA followed by Šídák’s multiple comparisons test (F), or unpaired t-test (G) (p< *0.05, **0.01, ***0.001, ****0.0001).

We hypothesized that E0771 growth is inhibited in lean mice relative to obese mice because lean adipocytes trigger ferroptosis *in vivo*, contributing to the lower tumor burden observed in lean mice (Fig. 1A) and with lean adipocyte co-injection (Fig. 1E). We confirmed that 4-HNE levels are elevated in the lysates of E0771 tumors from lean mice compared to obese mice (Fig. 2E, fig. S2K). Additionally, mice bearing E0771 tumors were treated with the ferroptosis inhibitor liproxstatin-1 (fig. S2L). Ferroptosis inhibition by liproxstatin-1 in tumors was confirmed with decreased levels of 4-HNE in treated tumors (fig. S2M). Strikingly, the growth and mass of E0771 tumors were dramatically increased in lean mice when ferroptosis was inhibited (Fig. 2F-H). In contrast, there was no change in tumor burden in the obese mice treated with liproxstatin-1 (fig. S2N). Thus, the lower tumor burden observed in lean mice is at least in part due to increased ferroptosis of breast cancer cells in this context.

### Lean adipocytes secrete ferroptosis inducer 9S-HODE

Next, we sought to identify the paracrine factor(s) secreted by lean, but not obese adipocytes that induce ferroptosis. To determine what type of secreted factor(s) causes this phenotype, we selectively depleted heat-labile proteins or lipids in the lean secretome using heat inactivation or hydrophobic compounds, respectively. Heat inactivation did not affect the ability of the lean secretome to suppress E0771 or SkBr3 cell growth (Fig. 3A, fig. S3A). In contrast, lipid depletion using two different methods, Charcoal Dextran or Bio-Beads SM-2, completely rescued growth in both cell lines (Fig. 3A, fig. S3A-B). Interestingly, lipid depletion using a third method, Cleanascite, did not rescue growth (Fig. 3A, fig. S1A-B), revealing that the three hydrophobic compounds have differential affinities for distinct lipid species. Leveraging these differential depletions, we next used an untargeted, global QTOF LC-MS lipidomics workflow to isolate the features responsible for the growth inhibition (Fig. 3B). Intriguingly, the features of interest had masses indicative of oxidized fatty acyls, with one significant hit corresponding to the formula C18H32O2, which a review of known annotations suggested to correspond to linoleic acid-derived oxylipins. To confirm our findings, we acquired a panel of linoleic acid-derived oxylipin standards and screened these against our samples using targeted QQQ LC-MS/MS. This definitively identified eight oxylipins in our samples (Fig. 3C). All eight oxylipins were enriched in the lean secretome, with 9-HODE exhibiting the greatest fold change compared to the obese secretome (Fig. 3C). Importantly, elevated levels of linoleic acid-derived oxylipins were observed in both lean female and male whole inguinal/mammary white adipose tissue compared to obese (Fig. 3D, fig. S3D). The extracted ion chromatograph confirmed the identity of 9-HODE with a characteristic fragmentation from *m/*z 295.2 -> 171.0 (fig. S3C).

**Fig. 3.**
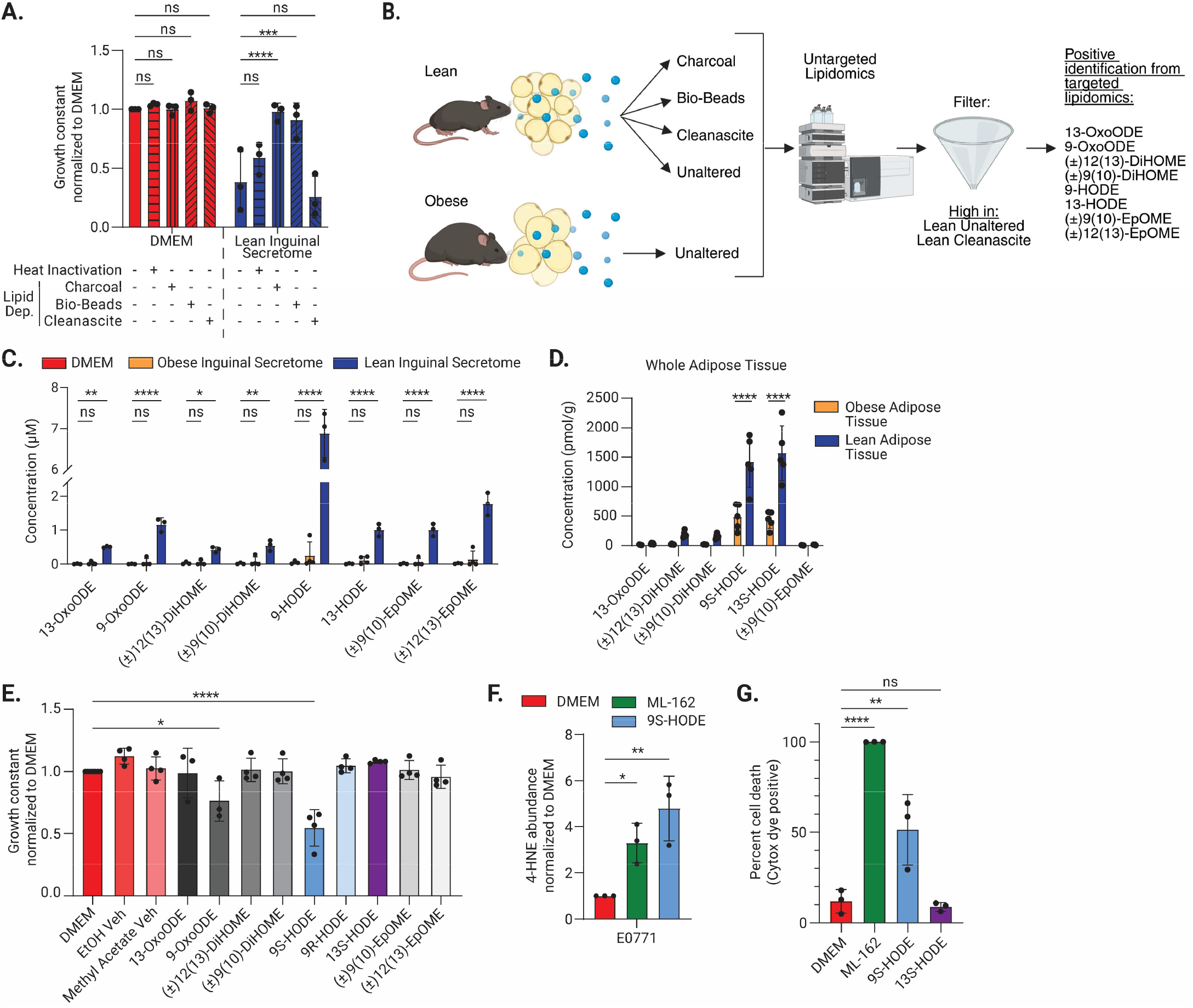
Lean inguinal adipocytes secrete ferroptosis inducer 9S-HODE. (**A**) Heat inactivation and lipid depletion of the lean inguinal adipocyte secretome. Lipid depletion using charcoal and Bio-Beads SM-2 Adsorbents rescues E0771 growth (n=3). (**B**) Schematic of the lipidomics methodology. (**C**) Relative concentrations of oxylipins identified in DMEM or three independent isolations of the obese or lean inguinal adipocyte secretome (n=3). (**D**) Relative quantification of oxylipins in lean or obese female mammary white adipose tissue (n=5). (**E**) Treatment of E0771 cells with individual lipid species supplemented into the culture media. 9S-HODE and 9-OxoODE are sufficient to inhibit E0771 breast cancer cell growth (n=4). (**F**) E0771 cells treated with 9S-HODE or ML-162 have increased levels of 4-HNE compared to DMEM control (n=3). (**G**) Cytotox dye incorporation confirms that ML-162 and 9S-HODE induce cell death, compared to DMEM control or 13S-HODE (n=3). (A-G) Data are mean ± SD; p-values calculated using two-way ANOVA followed by Dunnett’s multiple comparison test (A, C) or Šídák’s multiple comparisons test (D), or one-way ANOVA followed by Dunnett’s multiple comparison test (E-G) (p< *0.05, **0.01, ***0.001, ****0.0001).

To determine whether these oxylipins can suppress the growth of breast cancer cells, each lipid was individually supplemented to the standard culture media, and E0771 growth was assessed. Strikingly, 9S-HODE was sufficient to inhibit E0771 growth (Fig. 3E). Mild growth inhibition was also observed with 9-OxoODE, a metabolite of 9S-HODE (*36*), but not with any other lipid tested (Fig. 3E). Since our oxylipin analysis does not delineate chirality and 9S-HODE can occur in a racemic mixture with its enantiomer (*37*), the effect of 9R-HODE was also tested. 9R-HODE had no effect on E0771 growth (Fig. 3E). These data reveal a selective ability of 9S-HODE to inhibit E0771 growth. Relatively minor changes in the positioning or chirality of the hydroxyl group, as seen in 13S-HODE and 9R-HODE, completely eliminated the ability of this lipid to suppress breast cancer cell growth (Fig. 3E). Further, 9S-HODE was sufficient to induce ferroptosis (Fig. 3F) and promote cell death in E0771 cells (Fig. 3G). No cell death was observed with the highly related oxylipin 13S-HODE (Fig. 3G). Thus, lean adipocytes secrete the oxylipin 9S-HODE, which suppresses breast cancer cell growth by inducing ferroptosis.

### Lean adipocytes and 9S-HODE selectively suppress breast cancer cell growth

Given this strong breast cancer growth suppression, we next asked if 9S-HODE and the lean adipocyte secretome are also toxic to non-cancer cells. Intriguingly, little to no overall growth suppression was observed when two different normal mammary epithelial cell lines, NMuMG and EpH4, were treated with the lean secretome or 9S-HODE (Fig. 4A-B, fig. S4A). Similarly, no induction of ferroptosis was observed in these cells (fig. S4B). In fact, 9S-HODE was three times as potent at suppressing the growth of breast cancer cells than non-cancerous cells (fig. S4C). No significant difference was observed in cancer versus non-cancer cells treated with 13S-HODE (fig. S4D).

**Fig. 4.**
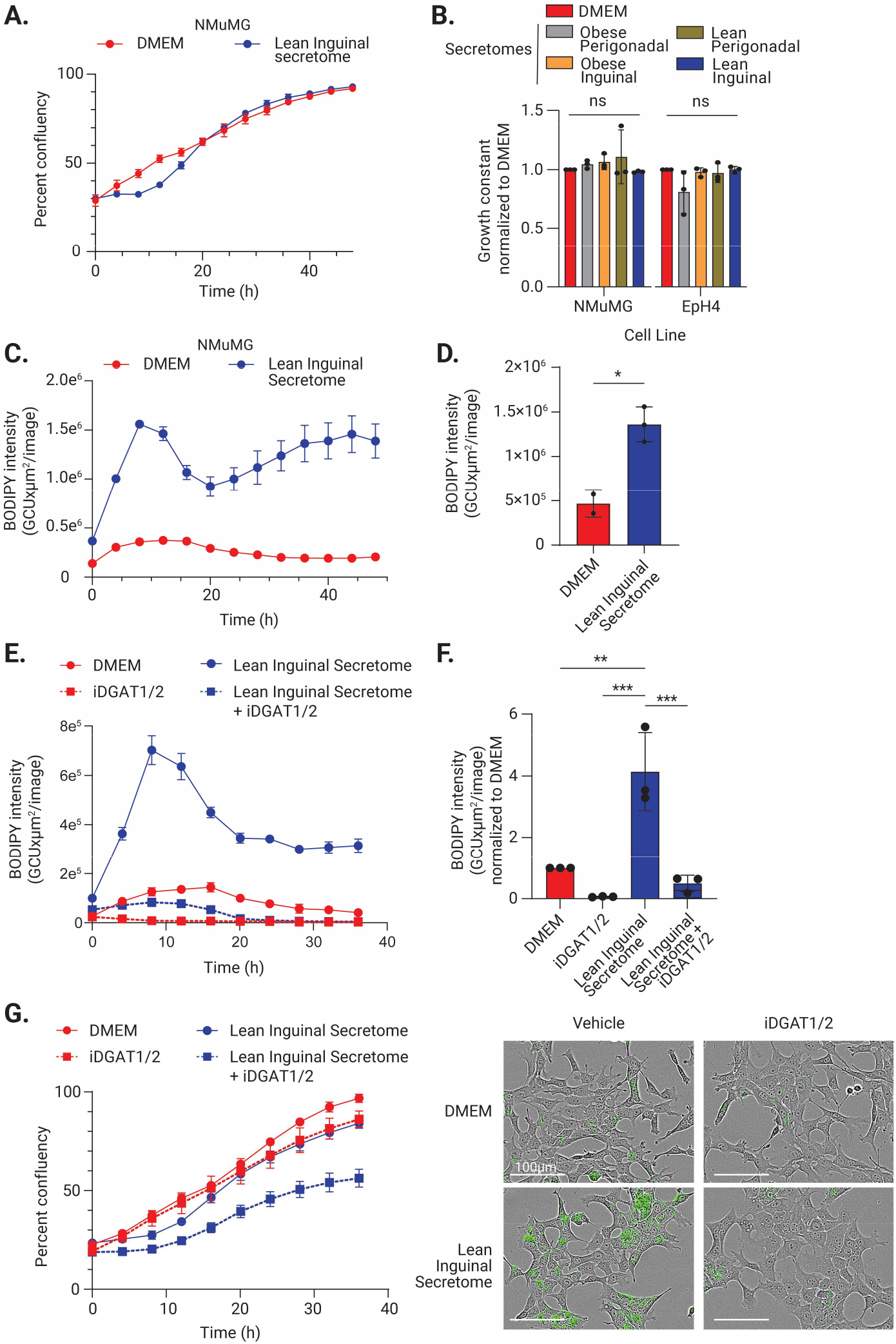
Normal mammary epithelial cells are protected from lean adipocytes. (**A**) Representative growth curve demonstrates that the lean inguinal adipocyte secretome induces a transient growth arrest in NMuMG breast epithelial cells. (**B**) Lean inguinal adipocyte secretome does not affect the overall growth constant of normal epithelial cell lines, NMuMG and EpH4 (n=3). (**C-D**) Lean inguinal secretome increases lipid droplet formation. (C) is representative of intracellular BODIPY intensity over time, (D) is quantification of BODIPY intensity 8 hours after addition of the lean inguinal secretome or DMEM control (n=3). (**E-F**) DGAT1/2 is required for epithelial cell protection from 9S-HODE. (E) is representative of BODIPY intensity over time, (F) is quantification of BODIPY intensity after 8 hours of treatment (n=3) with representative images below. (**G**) Representative growth curve for NMuMG cells treated with the lean inguinal adipocyte secretome, with or without iDGAT1/2. Treatment with iDGAT1/2 increases NMuMG cells’ sensitivity to the lean inguinal adipocyte secretome. (A, C, E, G) data are mean ± SD of technical replicates; (B-G) are mean ± SD of biological replicates; p-values calculated using one-way ANOVA followed by Dunnett’s multiple comparison test (B), unpaired t-test (D), or one-way ANOVA followed by Turkey’s multiple comparison test (F) (p< *0.05, **0.01, ***0.001).

Careful consideration of the growth kinetics revealed that non-cancerous epithelial cells underwent a transient cell cycle arrest immediately following the addition of the lean secretome (Fig. 4A). Lee et al. previously demonstrated that cells can avoid ferroptosis by transiently inducing a cell cycle arrest to allow for lipid droplet formation (*38*). To determine if non-cancerous epithelial cells form lipid droplets during the transient cell cycle arrest, we supplemented the media with a lipophilic fluorescent dye, BODIPY 493/503. Following the addition of the lean inguinal secretome, there was a dramatic spike in intracellular lipid droplets (Fig. 4C-D). DGAT1 and DGAT2 are required for lipid droplet formation (*39, 40*), and our gene expression analysis comparing E0771 and NMuMG cells showed high levels of *Dgat2* expression both basally and upon lean secretome treatment in NMuMG cells compared to E0771 cells (fig. S4E, data S1, S3). Inhibiting DGAT1 and DGAT2 (iDGAT1/2) completely prevented the formation of lipid droplets in NMuMG cells (Fig. 4E-F), and the loss of DGAT1/2-mediated lipid formation rendered NMuMG cells sensitive to growth inhibition by the lean secretome (Fig. 4G, fig. S4F) and 9S-HODE (fig. S4G-I). Thus, non-cancerous mammary epithelial cells can avoid cell death in response to 9S-HODE, presenting a possible therapeutic window to target breast cancer specifically.

### 9S-HODE possesses therapeutic potential

We hypothesized that loss of 9S-HODE with obesity could explain why obese adipocytes do not suppress breast cancer growth. To test whether 9S-HODE was sufficient to inhibit the growth of breast cancer cells *in vivo*, the obese secretome was supplemented with the same concentration of 9S-HODE as found in the lean secretome. Adding 9S-HODE was sufficient to suppress breast cancer growth (Fig. 5A). We next asked if 9S-HODE was sufficient to inhibit breast tumor growth *in vivo*. Obese and lean female C57BL/6J mice were injected with E0771 mammary tumors, and obese mice were randomized to receive intratumoral injections of vehicle or 9S-HODE three times a week once tumors reached enrollment size. Body weight was not affected by intra-tumoral injections (fig. S5A). Strikingly, 9S-HODE significantly inhibited tumor growth in obese mice (Fig. 5B-C, fig S5B). No effect on tumor growth was observed with 13S-HODE injections (fig. S5C-D), confirming the selective ability of 9S-HODE to inhibit breast cancer growth. We observed similar trends with another model, Py230 tumors (fig. S5E-G).

**Fig. 5.**
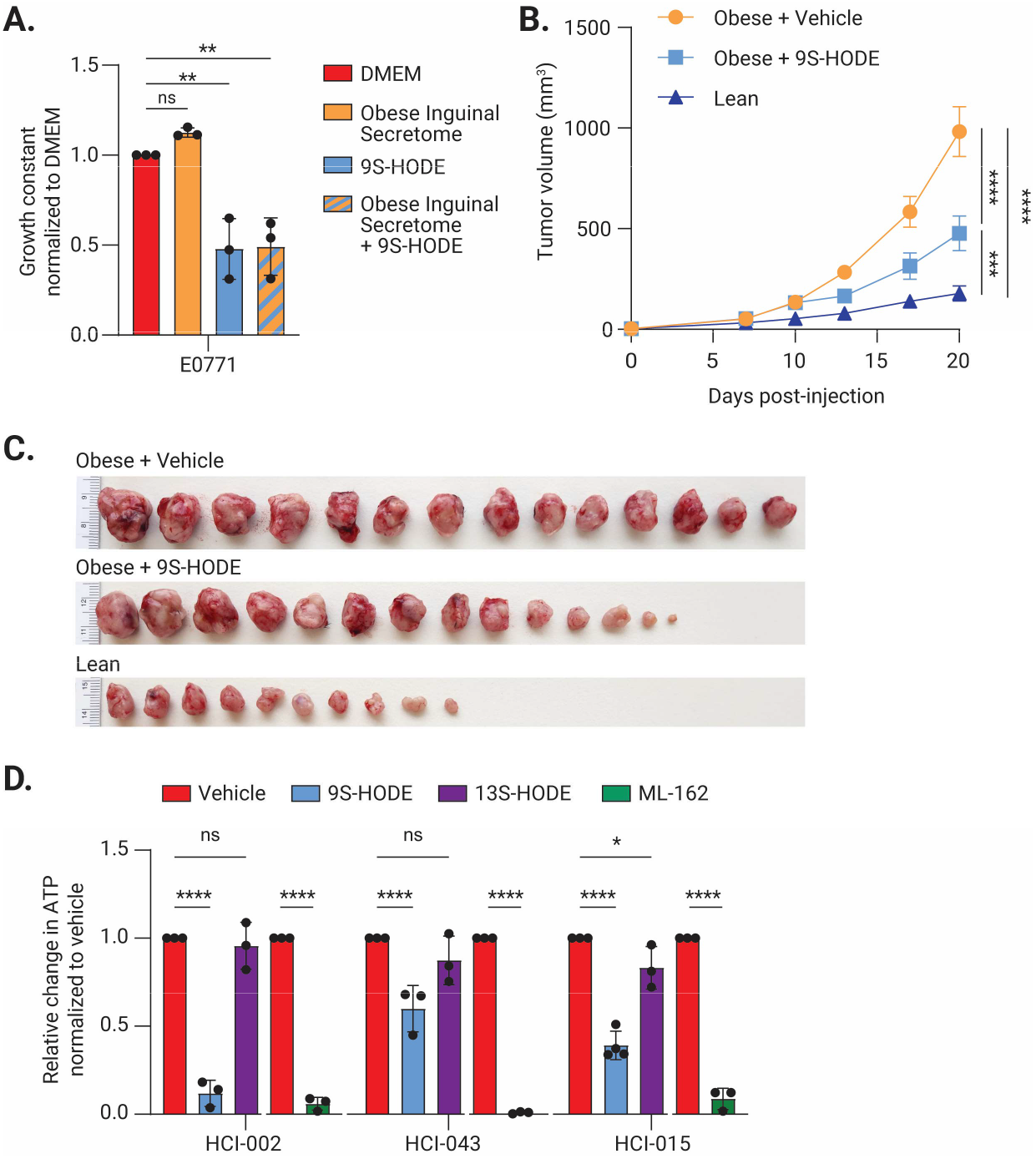
9S-HODE suppresses breast cancer growth. (**A**) The obese inguinal adipocyte secretome does not protect breast cancer cells against 9S-HODE. The growth constant for E0771 cells treated with the obese inguinal secretome of 3 individual isolations of mammary adipocytes (n=3) with or without 9S-HODE supplementation. (**B-C**) 9S-HODE intratumoral injection inhibits breast cancer tumor growth. (B) Growth of orthotopic E0771 mammary tumors in obese C57BL/6J mice treated three times a week with intratumoral injections of 9S-HODE (n=14) or vehicle (n=14) starting 10 days post-injection of breast cancer cells, or lean mice receiving no treatment (n=10). (C) are tumors at the experimental endpoint. (**D**) Breast cancer patient-derived xenograft organoids are sensitive to 9S-HODE treatment, quantified by Celltiter-Glo 3D, for three triple-negative patient-derived organoid lines. (A, D) Data are ± SD. (B) Data are ± SEM; p-values calculated using one-way ANOVA followed by Dunnett’s multiple comparison test (A, E) or two-way ANOVA followed by Turkey’s multiple comparison test (B) (p< *0.05, **0.01, ***0.001).

Finally, we sought to test if 9S-HODE showed tumor suppressive activity on breast cancer patient-derived organoids. Three different organoid lines, derived from patient-derived xenografts (*41*) were treated with 9S-HODE, 13S-HODE, or ML-162, and cell viability was measured using CellTiter-Glo. 9S-HODE, but not 13S-HODE is sufficient to inhibit the growth of patient-derived organoids (Fig. 5D). Together, these data demonstrate the therapeutic potential of 9S-HODE to reduce tumor growth.

## Discussion

We have identified a linoleic acid-derived oxylipin 9S-HODE that is secreted by lean mammary adipocytes and is sufficient to suppress breast cancer growth *in vitro* and *in vivo* by inducing ferroptosis. Obese adipocytes do not secrete 9S-HODE, which results in accelerated breast cancer growth in obese mice due to decreased levels of ferroptosis. Because 9S-HODE is a known human metabolite (*42*) and patient-derived breast cancer organoids are sensitive to 9S-HODE, we propose that breast cancer is accelerated in patients with obesity, at least in part, due to the loss of 9S-HODE. The discovery of 9S-HODE as a ferroptosis inducer also presents an exciting therapeutic opportunity. Our knowledge of endogenous ferroptosis inducers is limited to date. We show here that 9S-HODE potently kills both mouse and human breast cancer cells across all clinical subtypes, but spares non-cancerous breast epithelial cells. Together, these data suggest that lean adipocytes protect against breast cancer progression.

We observed a striking specificity of 9S-HODE, but not highly related lipids including 13S-HODE and 9R-HODE, to induce ferroptosis. We postulate the existence of a stereospecific regulator of ferroptosis in breast cancer cells. This may present an attractive therapeutic target in the substantial fraction of women diagnosed with breast cancer who are overweight or with obesity (*43*).

Most studies to date have focused on identifying tumor-promoting factors elevated in the tumor microenvironment of patients with obesity (*19-26*). To our knowledge, only one tumor-suppressive factor linked to lean adipose tissue, the adipokine adiponectin, has previously been described (*44*). Our work not only identifies another tumor-suppressive factor secreted by lean adipocytes, but demonstrates that breast cancer is accelerated in obese mice relative to lean mice in large part because breast cancer cells in lean mice are undergoing ferroptotic cell death. Few studies have characterized human breast cancer by obesity status, but a recent study similarly observed increased ferroptosis in breast cancer from lean patients compared to obese (*45*). There may be many more inhibitors of breast cancer that are secreted by lean adipose tissue and suppress growth by ferroptosis or other cellular mechanisms, but that have gone understudied due to an existing bias in the field to identify obesity-linked tumor-promoting mechanisms. The discovery of these anti-tumor factors in lean individuals, such as 9S-HODE, and the transcriptional and/or dietary mechanisms regulating their synthesis in lean versus obese adipocytes, may open up new therapeutic avenues to restore or augment their production in patients with obesity. Finally, we propose that beyond breast cancer, the lean tumor microenvironment may similarly protect against other types of cancer linked to obesity, such as colorectal cancer and endometrial cancer (*46*).

Obese inguinal or perigonadal adipocytes (lean or obese) do not inhibit breast cancer cell growth (n=3 independent adipocyte isolations). (**E**) Co-injection of lean inguinal adipocytes with MCF7 breast cancer cells decreases tumor growth compared to co-injection with obese adipocytes or a saline control (n=6). (A, E) Data are ± SEM, (B-D) data are mean ± SD. p-values calculated using two-way ANOVA followed by Šídák’s multiple comparisons test (A), or Turkey’s multiple comparisons test (B), or one-way ANOVA followed by Turkey’s multiple comparison test (C-E) (p< **0.01, ***0.001, ****0.0001).

## Acknowledgments

The authors acknowledge the use of Grammarly editing services and BioRender for schematic creation. We recognize the HSC Cell Imaging Core at the University of Utah for the use of the DeltaVision Ultra Widefield Microscope.

Lipidomics mass spectrometry analysis was performed at the Mass Spectrometry and Proteomics Core Facility at the University of Utah. The authors thank Dan Cuthbertson of Agilent Technologies for assistance in implementing iterative exclusion in the tandem mass spectrometry experiments. Mass spectrometry equipment was obtained through NCRR Shared Instrumentation Grant 1S10OD016232-01, 1S10OD018210-01A1, and 1S10OD021505-01. Research reported in this publication utilized the High-Throughput Genomics and Cancer Bioinformatics Shared Resource at Huntsman Cancer Institute at the University of Utah and was supported by the National Cancer Institute of the National Institutes of Health under Award Number P30CA042014. This project also utilized the Precision Cancer Models Shared Resource at Huntsman Cancer Institute at the University of Utah and was supported by the National Cancer Institute of the National Institutes of Health under Award Number P30CA042014. The content is solely the responsibility of the authors and does not necessarily represent the official views of the NIH.

## Funding

National Institutes of Health grant R01DK133455 (KIH) 5 For the Fight and Huntsman Cancer Institute (KIH)

V Foundation for Cancer Research (KIH) Pew Charitable Trusts (KIH)

University of Utah Philanthropic Group (KIH)

National Institutes of Health/National Cancer Institute grant U54CA224076 (ALW) Breast Cancer Research Foundation Founders Fund (ALW)

Department of Defense Breast Cancer Research Program Era of Hope Scholar Award W81XWH1210077 (ALW)

## Author contributions

Conceptualization: MCC, KIH Methodology: MCC, KIH, JAM Investigation: MCC, AEJ, EAB, JAM Visualization: MCC, JAM

Funding acquisition: KIH, JEC, ALW, DL Project administration: KIH, DL, JEC Supervision: KIH, DL, JEC, ALW Writing – original draft: MCC, JAM

Writing – review & editing: MCC, AEJ, EAB, KIH, ALW

## Data and materials availability

Sequencing data are available on NCBI GEO under accession number GSE300839. All remaining data needed to interpret the paper’s results are available in the main text or the supplementary materials.

## Supplementary Materials

Materials and Methods

Figs. S1 to S5

References (*47*–*60*)

Data S1-S3

