## Supplemental materials for "Lean breast adipocytes secrete an oxylipin that suppresses breast cancer via ferroptosis"

**The PDF file includes:**

Materials and Methods
Figs. S1 to S5
References (*47-60*)

**Other Supplementary Materials for this manuscript include the following:**

Data S1 to S3

### Materials and Methods

Cell Culture

Breast cancer cell lines (E0771, SKBR3, MCF7, BT474, MDA-MB-231, MDA-MB-468, Py230), and epithelial cell lines (NMuMG, EpH4) were purchased from American Type Culture Collection (ATCC). Breast cancer cell lines were cultured in Dulbecco's Modified Eagle medium (DMEM) supplemented with 10% Fetal bovine serum (FBS), 1% penicillin/streptomycin, and 1% glutamax. Py230 cells received additional supplementation of MITO+ Serum Extender (Corning® #355006) at a final concentration of 0.1%. Epithelial cell line NMuMG cells were cultured in DMEM with 10% FBS, 4.5 g/L glucose and 10 mcg/mL insulin. EpH4 cells were cultured in DMEM with 10% bovine calf serum (BCS) and 1.2 mcg/mL Puromycin. All cell lines were cultured at 37°C with 5% Co2. Patient-derived xenograft organoid lines (HCI-002, HCI-015, HCI-043) were cultured according to (*47*), maintained as established organoid lines, and then plated into 384-well plates for assaying. Treatments were added 24 hours after plating, and treatment response was measured 72 hours post-treatment using the CellTiter-Glo® 3D assay (*41*).

Animal Models

All manipulations were performed under the University of Utah, Institutional Animal Care and Use Committee–approved guidelines and policies. C57BL/6J (000664) mice were purchased from The Jackson Laboratory, age 3-6 weeks. Animals were housed at a constant temperature with a fixed 12-hour light/dark cycle with ad libitum food and water. Obese mice were generated by placing mice on High Fat Diet (Research Diets, D12492) at 6 weeks of age for a minimum of 9 weeks. Lean mice were maintained on a standard rodent chow diet (LabDiet 5053). Mice were age and gender matched and, within the same gender, were randomly assigned to treatment or control groups in all experiments and adipocyte isolations. Body composition was measured using NMR with the Bruker LF50.

Mammary Tumor Studies

For experiments with lean and obese mice, C57BL/6J female mice, age 15-18 weeks, were injected with 1 million E0771 breast cancer cells, resuspended in 30ul of Matrigel, into both the right and left fourth mammary fat pad. Lean only experiments used 8-week-old mice. Tumor volume and mouse weight were monitored twice a week for the course of the study. Tumor volume was calculated using ((length*width$^2$)/2) where the width was the smaller measurement, taken perpendicular to the length. For co-injection of MCF7 with primary adipocytes, mature adipocytes were isolated from 18-week-old lean and obese male C57BL/6J mice as previously described (*48*). 50 µL of packed adipocytes were gently mixed with 3 million MCF7 cells and resuspended using a cut pipette tip in 50 µL Matrigel. A small incision was made within the flank of 6-week-old NRG mice. A pipette tip, with the tip cut, was loaded with the adipocyte MCF7 solution, slid into the incision, and used to deposit the cells. Tumor volume was calculated as above. Mouse weight was monitored twice a week over the duration of the study. Liproxstatin-1 (Cayman Chemical 17730) experiments included daily intraperitoneal injections with liproxstatin-1 (10mg/kg) or vehicle for the entirety of the study beginning the day of primary E0771 tumor cell injection (*49, 50*). For intratumor injections of 100 µM 9S-HODE, 13S-HODE, or matched vehicle, were diluted in sterile saline. Mice were randomized to treatment groups when their tumors reached between 80-150mm$^3$, and were then treated 3 times a week for the duration of the study.

Adipocyte conditioned media

Adipocytes were isolated from lean and obese C57BL/6J mice (male mice age 15-25 weeks, female mice age 23-25 weeks) in accordance with the protocol outlined by Curtin and Jackson et al. (*48*). In brief, the adipose tissue was removed from the mice and minced until a homogeneous solution was reached. The tissue was then digested in a collagenase solution (100 mg of collagenase powder, 1 mL of 1 M HEPES buffer, 500 µL of 100x P188 (Sigma P5556), 50 µL of 1 M CaCl2 (Sigma C1016), 0.05 g of BSA (Sigma-Aldrich A7906), and 48.45 mL of Medium 199 (Sigma-Aldrich M4530)) for 10-12 min. The tissue was passed through a sieve and centrifuged. The adipocytes floated to the top and were then transferred to a new tube, where they were washed and packed. Post adipocyte isolation, the mature adipocytes were plated under trans well plates (Corning 3413) in DMEM + 10% FBS, 1% Pen-Strep, 1% Glutamax at 37 °C with 5% CO2. The media, containing secreted factors, was collected every 48 hours and passed thought a 0.44-micron filter to remove possible cellular debris. The secreted factor-filled media was aliquoted and immediately flash frozen in liquid nitrogen and stored at -80 °C. The adipocytes were then replenished with fresh media. Conditioned media collections lasted for 2 weeks. Secretomes used in this study were collected after 1 week in culture.

For heat inactivation experiments, the conditioned media was incubated at 100 °C for 10 min and then allowed to cool to room temperature. For lipid depletion experiments, the conditioned media was incubated with either charcoal dextran (Sigma C6241), cleanascite (Biotech support group X2555) or Bio-Beads SM-2 Adsorbents (Biorad 1528920). Charcoal Dextran was used at a ratio of 0.02g to 1mL, the conditioned media was incubated at 2 hours at room temperature, centrifuged at 2000g for 15 min, and then passed through a 0.44-micron filter. Cleanscite was used at a 1:4 ratio, the cleanascite beads were first washed once with DMEM, the conditioned media was then added and incubated at room temperature for 30 min, centrifuged at 2000 g for 15 minutes, and then passed through a 0.44-micron filter. Bio-Beads SM-2 were used at a ratio of 0.2g per 1 mL. Conditioned media was added and incubated for 2 hours at room temperature, centrifuges at 2000 g for 15 min, then passed through a 0.44-micron filter. For the growth assay experiments, the heat inactivated and lipid depleted conditioned media were mixed with fresh culture media in a 1:1 ratio. For lipidomics, the samples were submitted without dilution.

Growth Assay

Cells were plated into a 96-well plate with 5,000 to 10,000 cells per well in DMEM supplemented with 10% FBS, 1% penicillin/streptomycin, and 1% glutamax, and grown overnight. The next day, the media was changed and experimental conditions were added. For growth assays with secretome, the secretome-filled media was mixed 1:1 with fresh culture media. For treatment with lipids, 20 µM of lipids (unless otherwise noted) were supplemented into fresh culture media, then added to the cells. For experiments to induce ferroptosis, cells were treated with 1µM of ML-162 (XXX). For experiments including ferroptosis inhibitors, 10µM liproxstatin-1 (Cayman Chemical 17730) or 10µM ferrostatin-1 (Cayman Chemical 17729), the inhibitors were added after 24h. For experiments with Diacylglycerol acyltransferase (DGAT) inhibitors, 40 µM of DGAT1 (Sigma Aldrich SML0539) and 40 µM DGAT2 (Sigma Aldrich PZ0412) inhibitors, were added to the plate 6 h prior to the addition of secretome media or lipid treatment. After all treatments were added the plate was placed into the Incucyte Live Cell Imaging System (Sartorius) and imaged every 4 h for 72 h. The IncuCyte Zoom Analysis software was used to set up definitions to define the cell confluency, using phase images, and over time. To compare the growth of different cell lines with different growth rates, a growth constant was calculated. To calculate the growth constant, the doubling time for each condition was calculated in the exponential growth phase using Graphpad Prism to fit a nonlinear regression (curve fit). The growth constant was then calculated by dividing ln(2) by the doubling time.

To calculate the inhibitory concentrations for each lipid, the growth constant at specific concentrations was calculated and plotted vs the concentration of lipid. An inhibitory dose-response curve was calculated as [Inhibitor] vs. normalized response with variable slope using GraphPad Prism.

Immunofluorescent Imaging

Primary adipocytes were isolated from 16-week old C57BL/6 male mice, packed as described above, and then resuspended in 5 µg/mL CellMask (Invitrogen C10045) and 4.5 µg/mL Hoechst (Thermo Fisher Scientific 62249) in PBS and stained according to Curtin and Jackson et al. (JOVE) (*48*). In brief, following the completion of the staining, the adipocytes were diluted in 500µL of PBS, then transferred with a cut pipette tip to a microscope slide fitted with imaging spacers (Grace Biolabs 654006), and covered with a cover slip. The slides were imaged with a Nikon Spinning Disk Confocal.

RNA Sequencing Analysis

E0771, SkBr3 and NMuMG cells were plated at low confluency and allowed to grow for 36 hours, before treatment addition. The treatments, control DMEM or lean inguinal secretome was added to the cells for 16 hours. The cells were washed once with PBS then collected in RLT-bME buffer. The RNA was isolated using Qiagen RNeasy kit (74104).

RNA was sequenced using Illumina Sequencing with Agilent RNA ScreenTape Assay for QC and NovaSeq S4 Reagent Kit v1.5_150x150 bp Sequencing (2500 M read-pairs/lane). The library was prepped using NEBNext Ultra II Directional RNA Library Prep with poly(A) mRNA Isolation with adapter read 1 (AGATCGGAAGAGCACACGTCTGAACTCCAGTCA) and adapter read 2 (AGATCGGAAGAGCGTCGTGTAGGGAAAGAGTGT).

The mouse GRCm39 genome and gene annotation files were downloaded from Ensembl release 108 and a reference database was created using STAR version 2.7.9a (*51*). Optical duplicates were removed from the paired end FASTQ files using clumpify v38.34 and reads were trimmed of adapters using cutadapt 1.16 (*52*). The trimmed reads were aligned to the reference database using STAR in two pass mode to output a BAM file sorted by coordinates. Mapped reads were assigned to annotated genes using featureCounts version 1.6.3 (*53*). The output files from cutadapt, FastQC, FastQ Screen, Picard CollectRnaSeqMetrics, STAR and featureCounts were summarized using MultiQC to check for any sample outliers (*54*). Differentially expressed genes were identified using a 5% false discovery rate with DESeq2 version 1.40.2 (*55*). The log2 fold changes were shrunk using the normal shrinkage estimator and a ranked list of genes were analyzed using the fast gene set enrichment package (*56*). Pathways were analyzed using the fast gene set enrichment package (*56*), Ingenuity Pathway Analysis (*57*) and KEGG pathways from Enrichr. Pathways with less then 20 genes were excluded.

4-HNE ELISA

ELISA for 4-HNE was performed according to Abcam Lipid peroxidation Assay Kit instructions (ab238538). Briefly, wells were coated with 100uL of 4-HNE conjugate and incubated overnight at 4 °C. The next day, the wells were washed twice with PBS, and blocked with assay diluent for 1 hr. Samples and standards were loaded into the wells and incubated for 10 min at room temperature with shaking. The 4-HNE antibody was then added and incubated for 1 h at room temperature. The wells were then washed 3 times with wash buffer before the addition of the secondary antibody-HRP conjugate, that was then left to incubate for 1 h at room temperature with shaking. The wells were washed 3 times with wash buffer. The substrate solution was added at room temperature with shaking. The stop solution was added after 4-5 min and the absorbance read at 450nm.

Cytotox Dye and Caspase Cleavage

E0771 and SkBr3 cells were plated at sub confluency, after 24 hours the media was changed to include treatment such as standard growth media (DMEM), lean inguinal secretome, 10 μM ML-162, or 1μM Doxirubicin. Incucyte® Cytotox Dye was spiked in after 24 hours, Incucyte® Caspase-3/7 Dye was added concurrently with treatment. The Incucyte® Live-Cell Analysis Systems and software were used to analyze the accumulation of the active fluorescent substrate after 24-36 hours.

Cell Cycle Analysis

E0771 and SkBr3 cells were plated at sub confluency and grown for 36 hours. After 36hrs, the media was changed and treatments, Thymidine (2mM), Nocodazole (50ng/mL), Etoposide (10μM), lean inguinal secretome, or control media was added. After 16hrs, the cells were lifted, washed twice with PBS and fixed using ice cold 70% ethanol. Cells were then washed twice, permeabilized and stained with 0.15% Triton-X, 100ug/mL RNase (NEB T3018L), 50ug/mL Propidium Iodine (Invitrogen P1304MP) overnight. Cells were then washed and stained with Phospho-Histone H3 (Cell Signaling #3465) for 1 hour. Cells were then washed, resuspended in PBS and analyzed using a Canto fluorescent cell analyzer with software BD FACSDiva 6.1.3. Cell cycle analysis was performed using the Dean-Jett Fox univariate cell cycle module in FlowJo 10.7.1.

Immunoblot

Primary E0771 cell tumors from lean or obese mice were flash frozen, then ground, resuspended in radioimmunoprecipitation assay (RIPA) buffer (Thermo scientific 89900) containing protease inhibitor (Thermo Scientific, Pierce A32965), centrifuged at 12,000 g for 15 min and collected the supernatant in a new tube. Pierce BCA Protein Assay Kit was used to determine the protein content of each sample. An equal amount of each sample was combined with NuPAGE LDS Sample Buffer 4x (Invitrogen NP0007) and equal amounts of protein were loaded into a NuPAGE 4-12% Bis-Tris Gel (Invitrogen NP0323BOX). Mini Protein Gels were run in NuPAGE™ MES SDS Running Buffer (Thermo Fisher Scientific NP0002), followed by transfer onto nitrocellulose membranes (BIO RAD, 1620115) using Towbin Buffer (2.5 mM Tris, 19.2 mM glycine, pH 8.3) containing 20% methanol. Membranes were blocked with blocking buffer (3% BSA, TBST buffer (20 mM Tris, 150 mM NaCl, 0.1% Tween 20, pH 7.5)) for 30 minutes at room temperature, followed by incubation with primary antibody for 4-HNE (ab46545, 1:1000) in 1% BSA and in TBST buffer for 24 hours at 4ºC. The membrane was then washed 3 times in TBST buffer for 10 minutes and incubated with Immun-Star Goat Anti-Rabbit

(GAR)-HRP Conjugate (Bio-rad, 1705046) in 1% BSA, TBST for 1 h at room temperature. Membranes were washed 3 times in TBST buffer. Enhanced Chemiluminescence Substrate (Revity, NEL105001) was added to the membrane immediately before imaging and allowed to develop for 2-3 minutes. The membrane was then scanned on an Invitrogen iBright1500. The membrane was then washed two times in TBST buffer for 10 minutes and placed into primary β-actin (Cell Signaling 4967, 1:1000) or  in 1% milk in TBST for 16hrs at 4ºC. The membrane was then washed 3 times in TBST buffer for 10 minutes and incubated with IRDye secondary antibody (LiCor, 926-32212 1:20,000 or LiCor 1:20,000) in 1% milk, TBST for 1 hour at room temperature. The membrane was then washed 3 times in TBST buffer and scanned on an Odyssey CLx Imaging System (LI-COR).

Lipids

Lipids were purchased through Cayman Chemical in resuspended in ethanol or methyl acetate, 9R-HODE (38405), (±)12(13)-EpOME (52450), (±)9(10)-EpOME (52400), 13S-HODE (38610), 9S-HODE (38410), (±)9(10)-DiHOME (53400), 13S-OxoODE (38620), 9S-OxoODE (38420), (±)12(13)-DiHOME (10009832). Unless otherwise noted, 20µM of lipids were used for *in vitro* studies.

Chemicals

LC-MS-grade solvents and mobile phase modifiers, including methanol, acetonitrile, isopropanol, and formic acid, were obtained from Honeywell Burdick & Jackson (Morristown, NJ); methyl tert-butyl ether from Fisher Scientific (Waltham, MA); ammonium formate and ammonium acetate from Sigma–Aldrich/Fluka (St. Louis, MO); and phosphate-buffered saline from Life Technologies Corp. (Grand Island, NY). Lipid standards were sourced from Cayman Chemical (Ann Arbor, MI), including Linoleic Acid Oxylipins MaxSpec LC-MS Mixture (20794) and 9(S)-HODE-d4 (338410).

Untargeted Lipidomics

Sample Preparation: For untargeted lipidomics, 500 µL of conditioned media was transferred into 13 × 100 mm screw-capped glass test tubes with Teflon caps. In a randomized sequence, chloroform, methanol, and 10 µL of internal standard were added (8:4:3, v/v/v, chloroform/methanol/water). Samples were incubated on ice for 30 minutes with occasional vortexing, then centrifuged at 3,900 × g for 5 minutes at 4 °C. The organic (lower) layer was collected, while the aqueous (upper) layer was re-extracted with 2 mL of chloroform/methanol (2:1, v/v), vortexed briefly, and centrifuged again. The combined organic layers were evaporated to dryness under vacuum. Lipid extracts were reconstituted in 300 µL of isopropanol/acetonitrile/water (4:1:1, v/v/v) and transferred to LC-MS vials. A process blank and a pooled quality control (QC) sample were prepared concurrently.

LC-MS Analysis: Lipid separation was performed using an Acquity UPLC CSH C18 column (2.1 × 100 mm, 1.7 µm) with a VanGuard precolumn (5 × 2.1 mm, 1.7 µm) (Waters, Milford, MA) maintained at 65 °C. The system was connected to an Agilent HiP 1290 Sampler, Agilent 1290 Infinity pump, and Agilent 6545 Accurate Mass Q-TOF dual AJS-ESI mass spectrometer (Agilent Technologies, Santa Clara, CA). Samples were analyzed in a randomized order in both positive and negative ionization modes over an m/z range of 100–1700. For positive mode, the source gas temperature was 225 °C, drying gas flow 11 L/min, nebulizer pressure 40 psig, sheath gas temperature 350 °C, and sheath gas flow 11 L/min. The VCap voltage was 3500 V, nozzle voltage 500 V, fragmentor 110 V, skimmer 85 V, and octopole RF peak 750 V. For negative mode, the source gas temperature was 300 °C, drying gas flow 11 L/min, nebulizer pressure 30 psig, sheath gas temperature 350 °C, and sheath gas flow 11 L/min. The VCap voltage was 3500 V, nozzle voltage 75 V, fragmentor 175 V, skimmer 75 V, and octopole RF peak 750 V. Mobile phase A consisted of acetonitrile/water (60:40, v/v) with 10 mM ammonium formate and 0.1% formic acid, while mobile phase B consisted of isopropanol/acetonitrile/water (90:9:1, v/v/v) with 10 mM ammonium formate and 0.1% formic acid. For negative mode, ammonium acetate replaced ammonium formate. The chromatography gradient was as follows: 15% B at injection, increased to 30% B over 2.4 min, 48% B from 2.4–3.0 min, 82% B from 3.0–13.2 min, and 99% B from 13.2–13.8 min, held until 16.7 min, then returned to initial conditions and equilibrated for 5 min. The flow rate was 0.4 mL/min with injection volumes of 2 µL for positive and 8 µL for negative mode. Tandem mass spectrometry was performed using iterative exclusion with the same LC gradient at collision energies of 20 V (positive mode) and 27.5 V (negative mode). Lipid annotation was conducted using accurate mass and MS/MS matching with the Agilent Lipid Annotator library (*58*) and LipidMatch (*59*).

Targeted Oxylipin Assay

Sample Preparation: Lipid extraction was performed using a modified version of the method described by (*60*). Tissue samples were homogenized for 15 seconds in 1 mL of methanol/phosphate-buffered saline (MeOH/PBS, 1:9, v/v) and immediately placed on ice. The homogenates were then transferred to pre-chilled Eppendorf tubes containing MeOH/PBS (1:9, v/v) to achieve a final concentration of 30 mg/mL. Internal standards were added, followed by brief vortexing. Solid-phase extraction (SPE) was conducted using Phenomenex Strata-X reverse-phase polymeric SPE cartridges (30 mg/well; P/N: S300-191) on a Tecan ResolveX A200 platform (Tecan Group Ltd., Männedorf, Switzerland). The cartridges were conditioned with 3 mL of MeOH, followed by 3 mL of water. Samples were then loaded onto the activated cartridges and washed with 3 mL of MeOH/water (1:9, v/v). After discarding the waste, lipids were eluted with 1 mL of MeOH into a 96-well plate and evaporated to dryness under nitrogen. Dried lipid extracts were reconstituted in 150 µL of acetonitrile/isopropanol/water (4:1:1, v/v/v) and transferred to labeled LC-MS vials with micro-inserts. A process blank and a pooled quality control (QC) sample were prepared concurrently.

LC-MS Analysis: Oxylipins were separated on an Acquity UPLC CSH C18 column (2.1 × 50 mm, 1.7 µm) coupled to an Acquity UPLC CSH C18 VanGuard precolumn (5 × 2.1 mm, 1.7 µm) (Waters, Milford, MA) with the column temperature maintained at 40°C. Chromatographic separation was performed using an Agilent 1290 Infinity UPLC system, consisting of a HiP 1290 sampler and 1290 Infinity pump, coupled to an Agilent 6490 triple quadrupole (QqQ) mass spectrometer. Oxylipins were detected using dynamic multiple reaction monitoring (dMRM) in negative ion mode. The ion source parameters were as follows: gas temperature, 175°C; gas flow, 15 L/min; nebulizer pressure, 30 psi; sheath gas temperature, 250°C; sheath gas flow, 12 L/min. The capillary voltage was set to 3000 V, and the nozzle voltage was 1500 V. A 5 µL injection volume was used, and samples were analyzed in a randomized order. Pooled quality control (QC) samples were injected throughout the analytical sequence. The mobile phases consisted of (A) water with 0.1% formic acid and (B) acetonitrile with 0.1% formic acid. The chromatographic gradient was as follows: 25% B at 0 min, increased to 50% B at 2 min, ramped to 100% B at 15 min, held until 18 min, and then re-equilibrated to 25% B over 3 min. The flow rate was 0.4 mL/min. Analyte-specific transitions and retention times were determined using authentic standards from the Linoleic Acid Oxylipins MaxSpec LC-MS Mixture and supplemented with values reported in the literature (*60*). Quantification of oxylipins was performed using an isotopic dilution with the stable isotope-labeled internal standard by comparing peak area ratios of the target compounds to the internal standard.

<u>Statistical Analysis and Data Visualization</u>

Statistical parameters including the statistical test used, exact value of n, what n represents, and the distribution and deviation are reported in the figures and corresponding figure legends. Most data are represented as the mean ± standard deviation unless otherwise noted, data points show independent biological replicates, and the p-value was determined using unpaired two-tailed Student's t tests, one-way ANOVA, or two-way ANOVA and noted accordingly. Unless otherwise stated, statistical analyses were performed in GraphPad Prism.

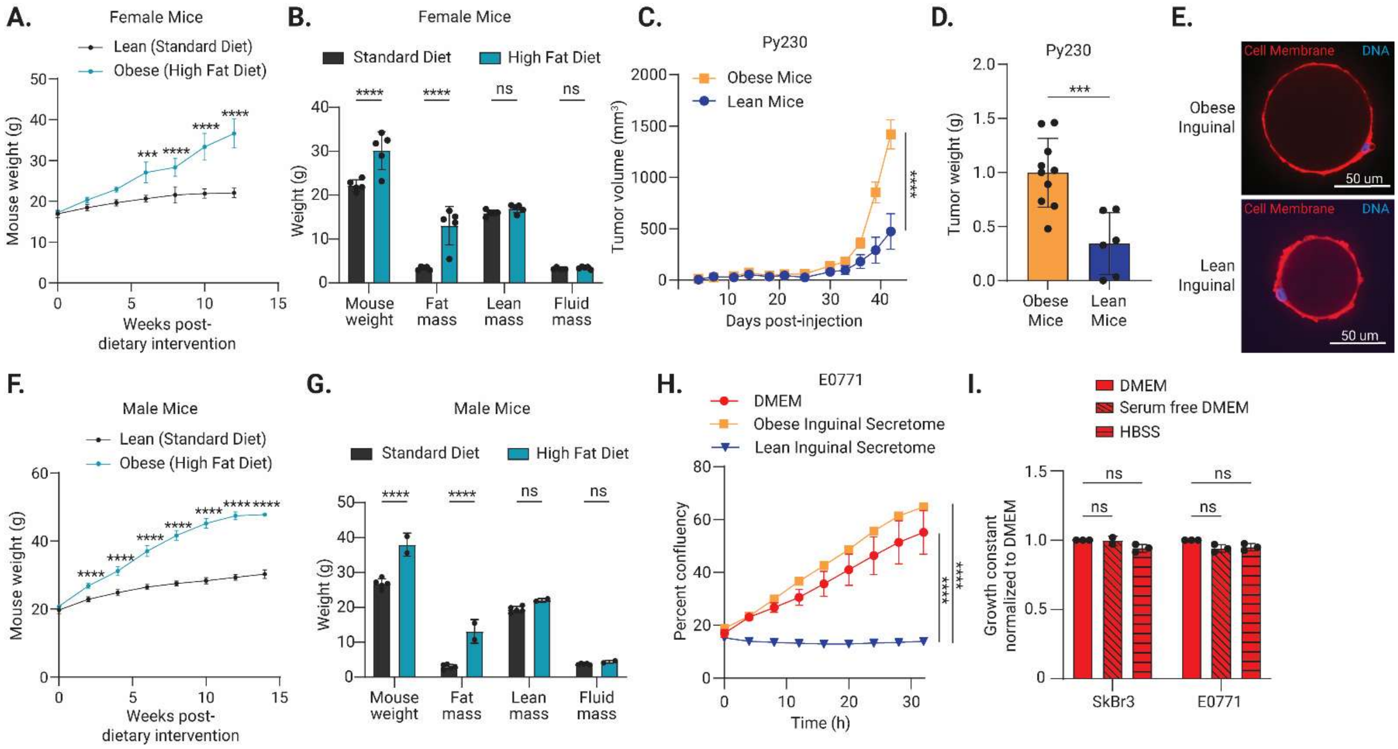


**Fig. S1. Lean inguinal adipocytes inhibit breast cancer cell growth.**

(**A-B**) (A) Body weight and (B) body composition of female C57BL/6J mice, after the start of dietary intervention at 6 weeks of age (n=5). (**C-D**) (C) Growth of orthotopic Py230 mammary tumors injected into C57BL/6 mice on a high fat (obese) (n=10) or normal chow (lean) diet over time and at (D) endpoint (n=6). (**E**) Representative images of isolated primary adipocytes from lean or obese inguinal fat pads, scale bar 50µm. (**F-G**) (F) Body weight and (G) body composition of male C57BL/6 mice, after the start of dietary intervention at 6 weeks of age (n=2-5). (**H**) Representative growth curve of E0771 cells treated with obese inguinal or lean inguinal adipocyte secretome collected from male mice. (**I**) Growth constant of SkBr3 or E0771 cells treated with 50% serum-free DMEM or HBSS, in place of secretome and 50% standard growth media (n=3). (A-B, D-I) Data are ± SD. (C) Data are ± SEM. P values calculated using two-way ANOVA followed by Šídák's multiple comparisons test (A-C F-G) or two-way ANOVA followed by Turkey's multiple comparison test (H) or one-way ANOVA followed by Dunnett's multiple comparison test (I) (p< *0.05, **0.01, ***0.001).

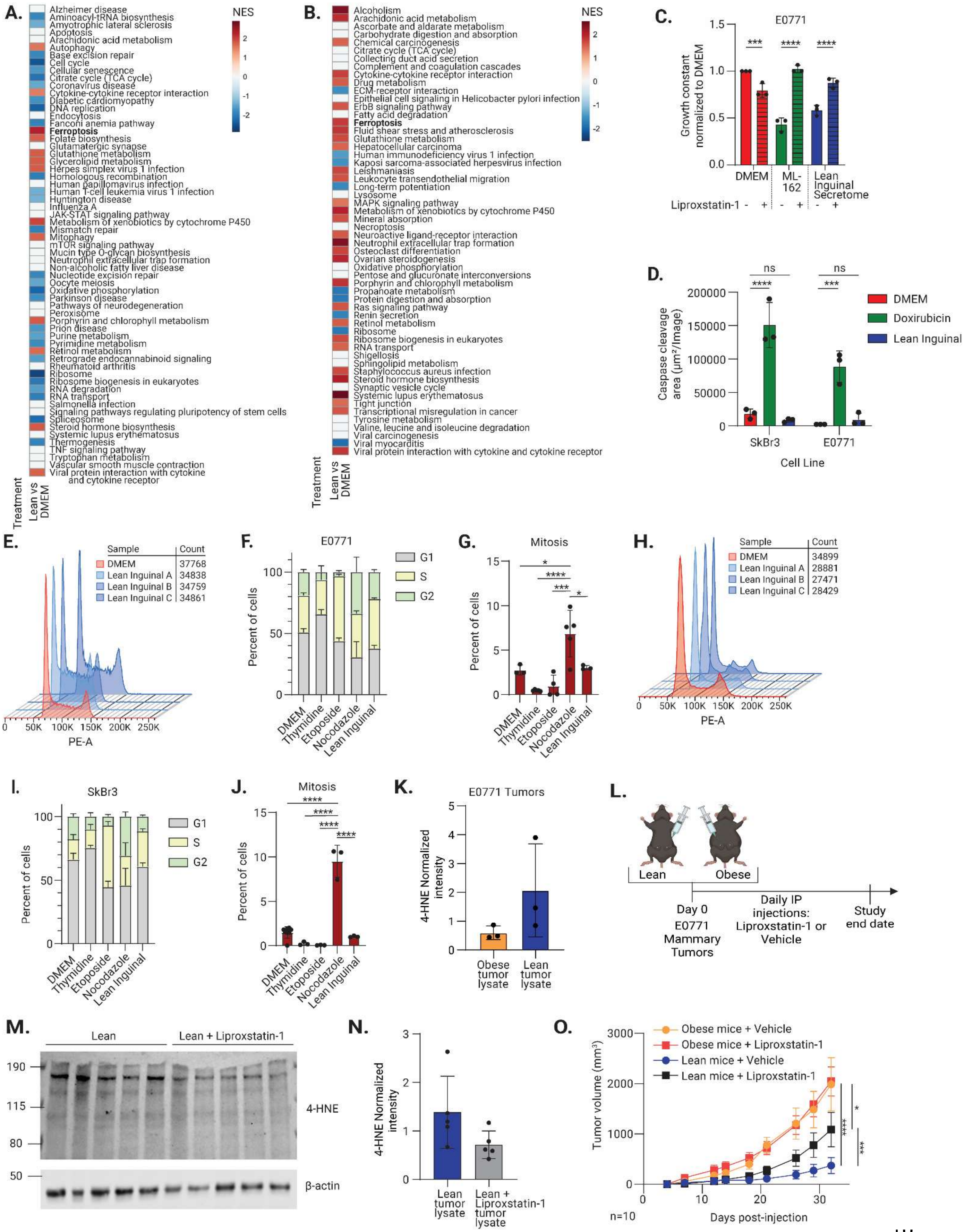
A.
NES
2
1
0
-1
-2
Alzheimer disease
Aminoacyl-tRNA biosynthesis
Amyotrophic lateral sclerosis
Apoptosis
Arachidonic acid metabolism
Autophagy
Base excision repair
Cell cycle
Cellular senescence
Citrate cycle (TCA cycle)
Coronavirus disease
Cytokine-cytokine receptor interaction
Diabetic cardiomyopathy
DNA replication
Endocytosis
Fanconi anemia pathway
Ferroptosis
Folate biosynthesis
Glutamatergic synapse
Glutathione metabolism
Glycerolipid metabolism
Herpes simplex virus 1 infection
Homologous recombination
Human papillomavirus infection
Human T-cell leukemia virus 1 infection
Huntington disease
Influenza A
JAK-STAT signaling pathway
Metabolism of xenobiotics by cytochrome P450
Mismatch repair
Mitophagy
mTOR signaling pathway
Mucin type O-glycan biosynthesis
Neutrophil extracellular trap formation
Non-alcoholic fatty liver disease
Nucleotide excision repair
Oocyte meiosis
Oxidative phosphorylation
Parkinson disease
Pathways of neurodegeneration
Peroxisome
Porphyrin and chlorophyll metabolism
Prion disease
Purine metabolism
Pyrimidine metabolism
Retinol metabolism
Retrograde endocannabinoid signaling
Rheumatoid arthritis
Ribosome
Ribosome biogenesis in eukaryotes
RNA degradation
RNA transport
Salmonella infection
Signaling pathways regulating pluripotency of stem cells
Spliceosome
Steroid hormone biosynthesis
Systemic lupus erythematosus
Thermogenesis
TNF signaling pathway
Tryptophan metabolism
Vascular smooth muscle contraction
Viral protein interaction with cytokine and cytokine receptor
Treatment
Lean vs DMEM
B.
NES
2
1
0
-1
-2
Alcoholism
Arachidonic acid metabolism
Ascorbate and aldarate metabolism
Carbohydrate digestion and absorption
Chemical carcinogenesis
Citrate cycle (TCA cycle)
Collecting duct acid secretion
Complement and coagulation cascades
Cytokine-cytokine receptor interaction
Drug metabolism
ECM-receptor interaction
Epithelial cell signaling in Helicobacter pylori infection
ErbB signaling pathway
Fatty acid degradation
Ferroptosis
Fluid shear stress and atherosclerosis
Glutathione metabolism
Hepatocellular carcinoma
Human immunodeficiency virus 1 infection
Kaposi sarcoma-associated herpesvirus infection
Leishmaniasis
Leukocyte transendothelial migration
Long-term potentiation
Lysosome
MAPK signaling pathway
Metabolism of xenobiotics by cytochrome P450
Mineral absorption
Necroptosis
Neuroactive ligand-receptor interaction
Neutrophil extracellular trap formation
Osteoclast differentiation
Ovarian steroidogenesis
Oxidative phosphorylation
Pentose and glucuronate interconversions
Porphyrin and chlorophyll metabolism
Propanoate metabolism
Protein digestion and absorption
Ras signaling pathway
Renin secretion
Retinol metabolism
Ribosome
Ribosome biogenesis in eukaryotes
RNA transport
Shigellosis
Sphingolipid metabolism
Staphylococcus aureus infection
Steroid hormone biosynthesis
Synaptic vesicle cycle
Systemic lupus erythematosus
Tight junction
Transcriptional misregulation in cancer
Tyrosine metabolism
Valine, leucine and isoleucine degradation
Viral carcinogenesis
Viral myocarditis
Viral protein interaction with cytokine and cytokine receptor
Treatment
Lean vs DMEM
C.
E0771
Growth constant normalized to DMEM
1.5
1.0
0.5
0.0
***
****
****
DMEM
ML-162
Lean Inguinal Secretome
Liproxstatin-1 - + - + - +
D.
ns
ns
****
***
Caspase cleavage area (µm²/Image)
200000
150000
100000
50000
0
DMEM
Doxirubicin
Lean Inguinal
SkBr3
E0771
Cell Line
E.
Sample | Count
DMEM | 37768
Lean Inguinal A | 34838
Lean Inguinal B | 34759
Lean Inguinal C | 34861
0 50K 100K 150K 200K 250K
PE-A
F.
E0771
Percent of cells
100
50
0
G1
S
G2
DMEM
Thymidine
Etoposide
Nocodazole
Lean Inguinal
G.
Mitosis
Percent of cells
15
10
5
0
*
****
***
*
DMEM
Thymidine
Etoposide
Nocodazole
Lean Inguinal
H.
Sample | Count
DMEM | 34899
Lean Inguinal A | 28881
Lean Inguinal B | 27471
Lean Inguinal C | 28429
0 50K 100K 150K 200K 250K
PE-A
I.
SkBr3
Percent of cells
100
50
0
G1
S
G2
DMEM
Thymidine
Etoposide
Nocodazole
Lean Inguinal
J.
Mitosis
Percent of cells
15
10
5
0
****
****
****
****
DMEM
Thymidine
Etoposide
Nocodazole
Lean Inguinal
K.
E0771 Tumors
4-HNE Normalized intensity
5
4
3
2
1
0
Obese tumor lysate
Lean tumor lysate
L.
Lean
Obese
Day 0
E0771 Mammary Tumors
Daily IP injections: Liproxstatin-1 or Vehicle
Study end date
M.
Lean
Lean + Liproxstatin-1
190
115
80
50
4-HNE
β-actin
N.
4-HNE Normalized intensity
3
2
1
0
Lean tumor lysate
Lean + Liproxstatin-1 tumor lysate
O.
Obese mice + Vehicle
Obese mice + Liproxstatin-1
Lean mice + Vehicle
Lean mice + Liproxstatin-1
Tumor volume (mm³)
3000
2000
1000
0
0
10
20
30
Days post-injection
n=10
****
*
***

**Fig. S2. Lean inguinal adipocytes induce ferroptosis.**

(**A-B**) RNA sequencing and pathway analysis using Enrichr of E0771 and SkBr3 cells, respectively, treated with DMEM control or lean inguinal adipocyte secretomes, pooled together from three different secretome collections, revealed an upregulation of ferroptosis. (**C**) Treatment with ferroptosis inhibitor Liproxstatin-1 rescues growth constant of E0771 cells treated with lean inguinal adipocyte secretome or ferroptosis inducer ML-162 (n=3). (**D**) SkBr3 and E0771 cells are not undergoing apoptosis as assessed by cleaved caspase 3/7 activity in cells treated with lean inguinal adipocyte secretome or doxorubicin (n=3). (**E-J**) Lean inguinal secretome induces an S-phase arrest in E0771 cells (E-G) and SkBr3 cells (H-J). (E, H) Whole histograms were obtained for single-cell events and cell-cycle histograms were constructed by selecting single-cell events with unfractionated DNA only. (F, I) Quantification of the cell-cycle histograms (n=3) (G, J) Percent of single cells events that are phospho-histone H3 (pH3) positive (n=3) (**K**) Quantification of immunoblot shown in Figure 2E. 4-HNE immunoblot bands from E0771 tumor lysates from lean or obese mice (n=3). (**L**) Schematic outlining the experiment. (**M-N**) Immunoblot of E0771 mammary tumor lysates from lean mice treated with or without liproxstatin-1 (Figure 2F), for levels of 4-HNE. (**O**) Growth of orthotopic E0771 mammary tumors injected into lean or obese C57BL/6 mice. Mice were then randomized to receive daily intraperitoneal injections of Liproxstatin-1 or vehicle (n=10). Data are ± SEM. (C-D, G-H, J-L) Data is mean ± SD. P values calculated using one-way ANOVA followed by Šídák's multiple comparisons test (C, D) or one-way ANOVA followed by Turkey's multiple comparisons test (G, J) (p< *0.05, **0.01, ***0.001, ****0.0001).

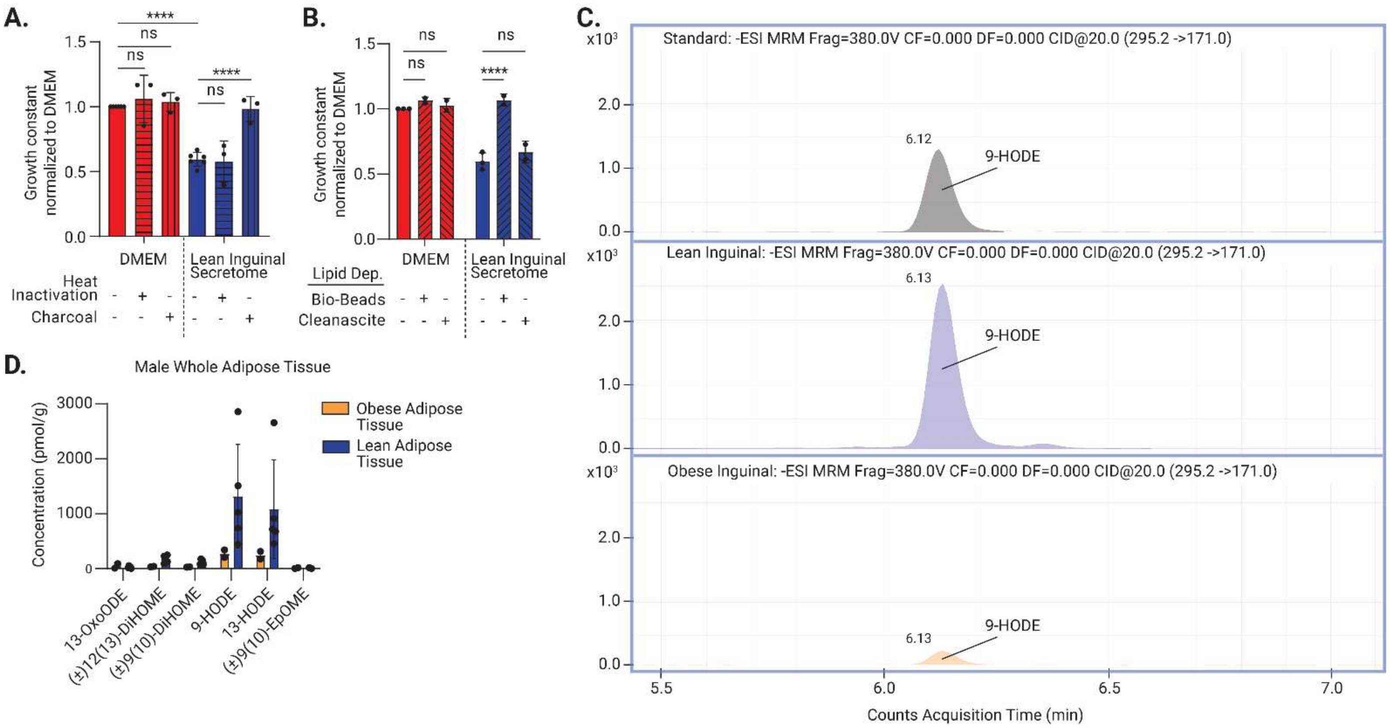


**Fig. S3. Lean inguinal adipocytes secrete 9S-HODE.**

(**A-B**) Heat inactivation and lipid depletion using charcoal dextran, Bio-Beads SM-2 Adsorbents, or Cleanascite of the lean inguinal adipocyte secretome. The lean inguinal adipocyte secretome with or without treatment was added to SkBr3 cells. (A) Three independent replicates, (B) three technical replicates. (**C**) Representative extracted ion chromatograph of oxylipin standard, lean inguinal adipocyte secretome, and obese inguinal adipocyte secretome samples highlighting relative 9-HODE abundance in the lean inguinal adipocyte secretome compared to the obese inguinal adipocyte secretome. MRM fragmentation 380.0V with transition 295.2->171.0. (**D**) Quantification of oxylipins in male inguinal white adipose tissue (n=3). (A-B, D) Data are mean ± SD. P values calculated using one-way ANOVA followed by Šídák's multiple comparisons test (A-B) (p< ****0.0001).

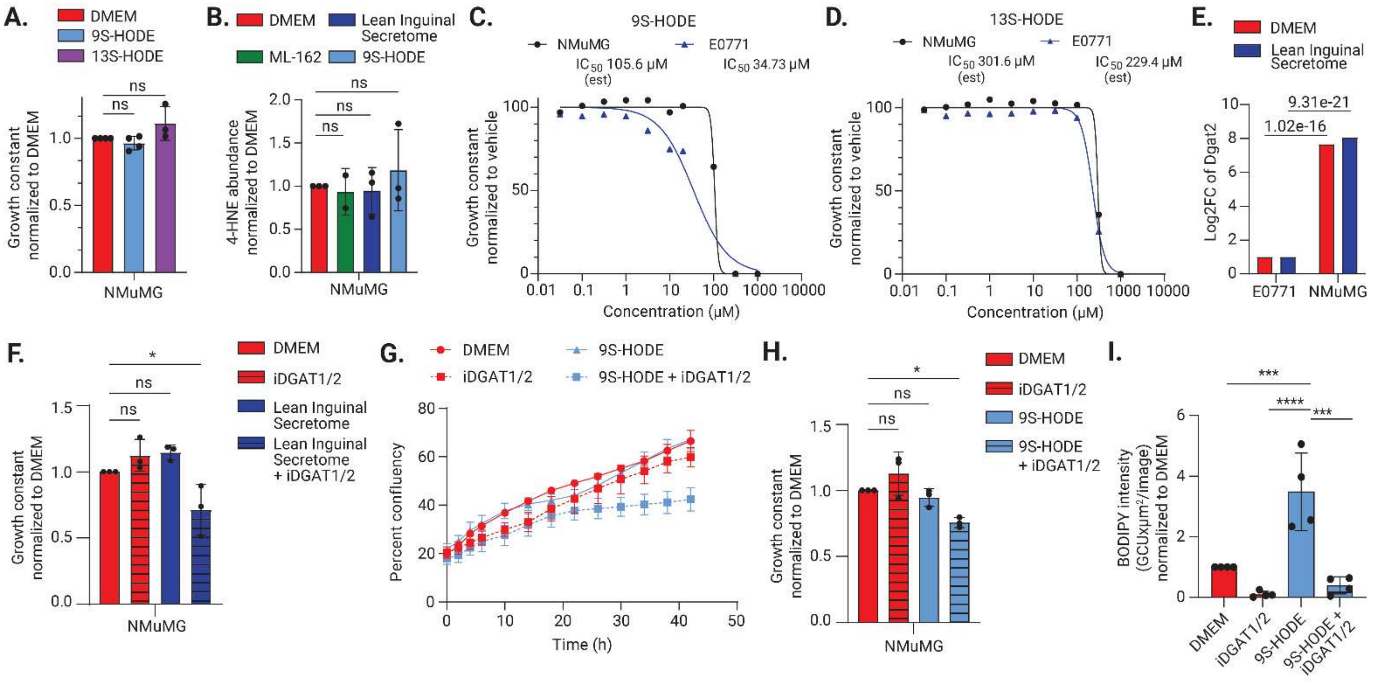


**Fig. S4. Normal mammary epithelial cells are protected 9S-HODE.**
(**A**) Growth constant of NMuMG cells treated with 9S-HODE or 13S-HODE (n=3). (**B**) Measure of 4-HNE abundance in cell lysates of NMuMG cells with ML-162, lean inguinal adipocyte secretome or 9S-HODE by 4-HNE ELISA (n=3). (**C-D**) $IC_{50}$ of 9S-HODE on E0771 and NMuMG cells, 34.73µM and 105.6µM (est) respectively. The estimated IC50 of 13S-HODE on E0771 and NMuMG cells is 229.4µM and 301.6µM, respectively. (**E**) Log2 fold change of Dgat2 levels in E0771 and NMuMG cells pre- and post-treatment with lean inguinal adipocyte secretome, quantified from Data S1 and S3. (**F**) Growth constant for NMuMG cells treated with lean inguinal secretome with iDGAT1/2 treatment (n=3). (**G-H**) (G) Representative growth curve and (H) growth constant of NMuMG cells treated with 9S-HODE with or without iDGAT1/2. (n=3). (**I**) BODIPY intensity as a marker of lipid droplet formation in NMuMG cells treated with 9S-HODE with or without iDGAT1/2, (n=3). (A-B, E-F, H-I) Data are mean ± SD. P values calculated using one-way ANOVA followed by Dunnett's multiple comparison test (A-B, F-H) or one-way ANOVA followed by Turkey's multiple comparisons test (F) ($p<$ *0.05, **0.01, ***0.001, ****0.0001).

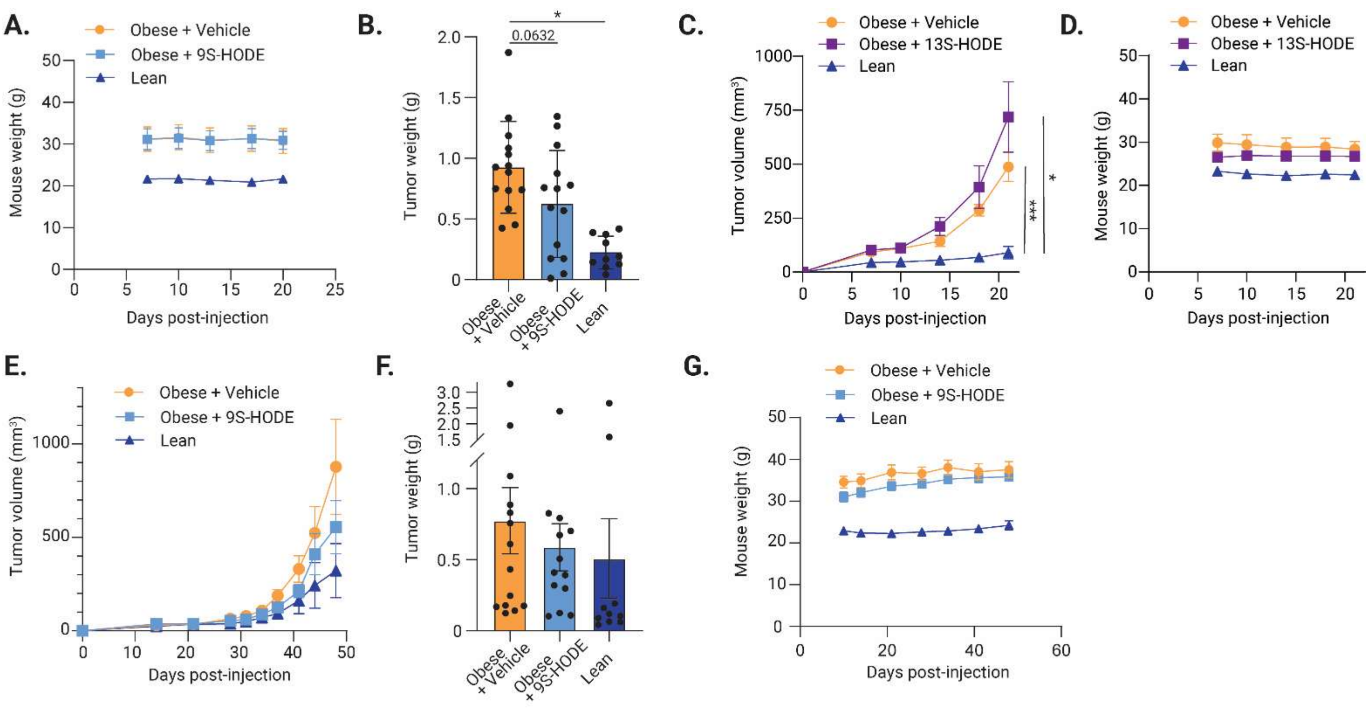


**Fig. S5. 9S-HODE suppresses breast cancer growth.**

(**A**) Body weight of mice over time, post-injection with E0771 tumor cells and throughout intratumoral treatment with 9S-HODE or vehicle, as in Figure 5B (n=7 obese + vehicle, n=7 obese + 9S-HODE, n=5 lean) (**B**) Tumor weights of E0771 tumors at endpoint from Figure 5B. Data is mean ± SD. (**C**) Tumor volume of E0771 mammary tumors in obese C57BL/6 mice treated with an intratumor administration of either vehicle (n=6) or 13S-HODE (n=6) (**D**) Body weight of mice over time, post-injection with E0771 tumor cells and throughout intratumoral treatment with 13S-HODE or vehicle, as in S5C (n=3 obese + vehicle, n=3 obese + 13S-HODE). (**E**) Growth of orthotopic Py230 mammary tumors in obese C57BL/6J mice treated with intratumoral injections of 9S-HODE (n=14) or vehicle (n=13), lean mice received no treatment (n=10). (**F**) Tumor weights of Py230 tumors at endpoint from figure S5E. (**G**) Body weight of mice over time, post-injection with Py230 tumor cells and throughout intratumoral treatment with 9S-HODE or vehicle, as in fig S5E (n=7 obese + vehicle, n=7 obese + 9S-HODE, n=5 lean). (A-G) Data are mean ± SEM. P values calculated using one-way ANOVA followed by Dunnett's multiple comparison test (B) or two-way ANOVA followed by Turkey's multiple comparisons test (C) (p< *0.05, **0.01, ***0.001, ****0.0001).

**Data S1. Significant gene changes in E0771 cells by lean inguinal secretome (separate file)**

Results of RNA sequencing showing genes that are upregulated or downregulated in E0771 breast cancer cells treated with the lean inguinal adipocyte secretome compared to standard culture media. Results are pooled from 3 independent treatments with different lean inguinal adipocyte secretomes.

**Data S2. Significant gene changes in SkBr3 cells by lean inguinal secretome (separate file)**

Results of RNA sequencing showing genes that are upregulated or downregulated in SkBr3 breast cancer cells treated with the lean inguinal adipocyte secretome compared to standard culture media. Results are pooled from 3 independent treatments with different lean inguinal adipocyte secretomes.

**Data S3. Significant gene changes in NMuMG cells by lean inguinal secretome (separate file)**

Results of RNA sequencing showing genes that are upregulated or downregulated in NMuMG breast cancer cells treated with the lean inguinal adipocyte secretome compared to standard culture media. Results are pooled from 3 independent treatments with different lean inguinal adipocyte secretomes.
